# Reactive or transgenic increase in microglial TYROBP reveals a TREM2-independent TYROBP-APOE link in wild-type and Alzheimer’s-related mice

**DOI:** 10.1101/2020.08.18.254649

**Authors:** Mickael Audrain, Jean-Vianney Haure-Mirande, Justyna Mleczko, Minghui Wang, Jennifer K. Griffin, Paul Fraser, Bin Zhang, Sam Gandy, Michelle E. Ehrlich

## Abstract

Microglial TYROBP (also known as DAP12) has been identified by computational transcriptomics as a network hub and driver in late-onset sporadic Alzheimer’s disease (AD) and as an important regulator of the microglial environmental sensing function. TYROBP is the transmembrane adaptor of AD-related receptors TREM2 and CR3, but importantly, TYROBP interacts with many other receptors, and little is known about its roles in microglial action and/or in the pathogenesis of AD. Herein, using dual RNA *in situ* hybridization and immunohistochemistry, we demonstrate that endogenous *Tyrobp* transcription is increased specifically in recruited microglia in the brains of wild-type and AD-related mouse models. To determine whether chronically elevated TYROBP might modify microglial phenotype and/or progression of AD pathogenesis, we generated a novel transgenic mouse overexpressing TYROBP in microglia. TYROBP-overexpressing mice were crossed with either *APP/PSEN1* or *MAPT^P301S^* mice, resulting in a decrease of the amyloid burden in the former and an increase of TAU phosphorylation in the latter. Apolipoprotein E (*Apoe*) transcription was upregulated in *MAPT^P301S^* mice overexpressing TYROBP and transcription of genes previously associated with *Apoe*, including *Axl*, *Ccl2*, *Tgfβ* and *Il6*, was altered in both *APP/PSEN1* and *MAPT^P301S^* mice overexpressing TYROBP. Lastly, *Tyrobp* and *Apoe* mRNAs were clearly increased in *Trem2*-null mice in microglia recruited around a cortical stab injury or amyloid-β (Aβ) deposits. Conversely, microglial *Apoe* transcription was dramatically diminished when *Tyrobp* was absent. Our results provide compelling evidence that TYROBP-APOE signaling in the microglial sensome does not require TREM2. We propose that activation of a TREM2-independent TYROBP-APOE signaling could be an early or even initiating step in the transformation of microglia from the homeostatic phenotype to the Disease-Associated Microglia (DAM) phenotype.

## INTRODUCTION

Microglia play a sentinel role in the brain, capable of detecting a wide variety of environmental stimuli, including microbial pathogens, aggregated proteins (such as amyloid-β; Aβ) and cellular debris (such as membrane fragments). This sensing activity is an essential part of the host response and is broad in its scope, sometimes triggering homeostatic adjustment, while, at other times, host defense. Microglia are also of interest in neurodegenerative diseases due to proteinopathies, e.g. Alzheimer’s disease (AD), in which large genetic studies have reported increased disease risk linked to many loci associated with microglial genes implicated in clearance of Aβ peptides^1–7^. More recently, transcriptomic analyses have revealed distinct profiles and signatures for microglia associated with aging and age-related diseases, indicating that a wide range of specific proteins in microglia underlies sensing, activation, and/or other cellular responses. Using RNA sequencing, Hickman *et al.* (2013) identified 100 transcripts highly enriched in microglia and coined the term “sensome” to describe this class of microglial transcripts^8^. Network analysis of this list identified a *TYROBP* (for tyrosine kinase binding protein; also known as *DAP12*, for DNAX activating protein-12)-centered pathway with 24 of these 100 genes interacting directly with *TYROBP* and 20 interacting indirectly with *TYROBP*. Concurrently, members of our team employed an integrative network-based approach and identified *TYROBP* as a key network driver in sporadic late-onset Alzheimer’s disease (AD)^9^. More recently, Keren-Shaul *et al*.^10^ used single-cell RNA sequencing in mouse to define a specific microglial phenotype that they termed “Disease-Associated Microglia” (DAM). *Tyrobp* was one of the genes most robustly upregulated in the proposed earliest stage of transition of microglia from the basal “homeostatic state” into the DAM phenotype.

TYROBP is a transmembrane signaling polypeptide that contains an immunoreceptor phospho-tyrosine-based activation motif (ITAM) in its cytoplasmic domain. TYROBP is expressed in microglia and serves as an adaptor for a variety of immune receptors, including two molecules closely linked to AD pathogenesis: TREM2 (triggering receptor expressed on myeloid cells 2) and CR3 (complement receptor 3). TREM2 is expressed at the plasma membrane of microglia in the brain and some mutations and polymorphisms of *TREM2* are linked to autosomal dominant AD or sporadic late-onset AD^11^. Other *TREM2* mutations can also cause a polycystic leukoencephalopathy osteodystrophy also known by the eponym Nasu-Hakola disease^12^. Most *TYROBP* mutations represent loss-of-function mutations and also result in Nasu-Hakola disease^13^. Similarly, TYROBP genetic variants have also been identified in early-onset AD^14^.

TREM2 is required in order for microglia to limit growth of amyloid-β (Aβ) deposits and is essential for a full transition of homeostatic microglia to a DAM state. Keren-Shaul *et al*.^10^described a two-stage program for DAM transition with a *Trem2*-independent step (stage 1) where *Tyrobp* and other genes are upregulated, followed by a *Trem2*-dependent step during which both *Tyrobp* and *Trem2* are upregulated (stage 2). Krasemann *et al.*^15^described a very similar microglial signature associated with neurodegenerative diseases, designated the “MGnD” phenotype, and showed that the transition from homeostatic to MGnD microglia was both TREM2- and APOE-dependent with a TREM2-APOE signaling pathway driving the transition from homeostatic microglia to MGnD. Notably, when DAM and MGnD are compared, Keren-Shaul *et al*^10^. also observed by single-cell RNA sequencing an apparent sequence of events whereby *Tyrobp* was upregulated prior to the upregulation of *Trem2*. For clarity, since DAM and MGnD microglia appear to share key features of the phenomena described here, we will refer only to DAM for the remainder of this report. Insofar as we are aware, principles established here underpin both DAM and MGnD.

In light of the central role of TYROBP in the microglial sensome^8^, its key role as adaptor for multiple microglial receptors^16^, its upregulation in the early *Trem2*-independent DAM stage 1^10^, and its upregulation in AD^9^, we employed a number of strategies aimed at interrogation of the causes and consequences of *TYROBP* upregulation. Using dual RNA *in situ* hybridization and immunohistochemistry, we found that *Tyrobp* mRNA level is significantly increased when microglia are recruited, including in wild-type (WT) mouse, in an *APP/PSEN1* transgenic mouse model of cerebral Aβ amyloidosis, and in a *MAPT^P301S^* transgenic mouse model of tauopathy. To determine whether elevated TYROBP can modify microglial phenotype and AD pathogenesis, we generated a novel transgenic mouse, designated the *Iba1^Tyrobp^*mouse, wherein the *Iba1* promoter was used to drive overexpression of a mouse *Tyrobp* transgene in microglia. The *APP/PSEN1* and *MAPT^P301S^* phenotypes were both altered in the setting of a constitutive increase in TYROBP, as was the transcription of *Apoe* and associated genes. Finally, using two mouse models of cerebral Aβ amyloidosis and a mouse model of penetrating cortical stab injury, we show that upregulation of *Tyrobp* and *Apoe* does not require *Trem2*, but that upregulation of microglial *Apoe* requires *Tyrobp* to reach normal levels.

Our results provide compelling evidence that TYROBP-APOE signaling in the microglial sensome is independent of *Trem2*, and we propose that activation of this pathway could be an early or even initiating step in the transformation of microglia from the homeostatic phenotype to the DAM phenotype.

## MATERIALS & METHODS

### Animals

All experimental procedures were conducted in accordance with the NIH Guidelines for Animal Research and were approved by the Institutional Animal Care and Use Committee (IACUC) at Icahn School of Medicine at Mount Sinai. TYROBP-overexpressing mice, named after *Iba1^Tyrobp^* mice, were constructed by cloning the mouse *Tyrobp* and *Egfp* sequences separated by an internal ribosome entry site (IRES) under the mouse *Iba1* promoter in a pBluescript II SK(+) vector^17^. GFP protein was not detected by immunohistochemistry. Microinjection were performed in C57BL/6J mice by Dr Kevin Kelly here at Mount Sinai. *APP^KM670/671NL^/PSEN1^∆exon9^* (*APP/PSEN1*) and *MAPT^P301S^* (PS19) mice were obtained from Jackson Laboratories. *Tyrobp* knockout (*Tyrobp^−/−^*) mice were obtained from Taconic/Merck Laboratory. *TgCRND8* were obtained from Dr Paul Fraser. *Trem2* knockout (*Trem2^-/^*^-^) mice were constructed by use of targeted homologous recombination to remove exons 1 and 2, thereby deleting the start codon and the major extracellular IgG domain. In contrast to the reported Velocigene construct, the direction of the Hygromycin cassette was “reversed”. Crucially, in agreement with two other models, but in contrast to the Velocigene construct, RT-qPCR analyses confirmed (data not shown) specific loss of Trem2 expression without the perturbation of Treml1 expression observed in the Velocigene construct^18^. For the penetrating cortical stab-injured mice, WT, *Tyrobp^−/−^* and *Trem2^−/−^* mice were anesthetized by an intraperitoneal injection of ketamine/xylazine (80/16 mg/kg body weight) and placed in a stereotactic frame (Stoelting, Wood Dale, IL, USA). The skull was drilled and an empty Hamilton syringe was introduced for 1 minute at the following coordinates (relative to bregma): anteroposterior, −0.3 mm; mediolateral, +3 mm; dorsoventral, −2mm. Mice were killed 3 days after.

### Tissue collection and sample preparation

Mice were anesthetized in a CO_2_ chamber then transcardially perfused with 20 ml ice-cold phosphate buffered saline (PBS). One hemisphere was post-fixed by incubation for 48 h in 4% paraformaldehyde (PFA) and cut into 35 μm sections with a vibratome (Leica) for histological analyses (brains used for RNAscope were perfused with both PBS and 4% PFA prior post-fixation and cut in 10 μm sections with a cryostat). The contralateral hemisphere was dissected for isolation of the hippocampus and cortex. Hippocampus was used for RT-qPCR and RNA sequencing, whereas cortical samples were homogenized in a RIPA buffer (Thermo Scientific, #89901) containing phosphatase and protease inhibitors (Thermo Scientific, #78442), centrifuged for 20 min at 15,000 × g and the supernatant was used.

### Immunohistochemistry

Sections were washed with 0.1% Triton in PBS, non-specific binding was eliminated by incubation with 0.1% Triton in PBS/5% goat serum, and then each slice was incubated with one of a panel of primary antibodies as follows: AT8 anti-pTAU pSer202/Thr205 (1:1000, Thermo Scientific, #MN1020); anti-pTAU Thr205 (1:1000, Invitrogen, #44-738G), anti-IBA1 (1:1000, Wako, #019-19741), anti-CD68 (1:500, Bio-Rad, #MCA1957); anti-6E10 (1:1000, Covance, #9320500), anti-TYROBP (1:500, antibody provided in collaboration with Richard W. Cho at Cell Signaling Technology). For fluorescent immunostaining, sections were incubated with the appropriate secondary antibody: anti-rabbit IgG Alexa Fluor 488 or 568 (1:2000, Invitrogen); anti-mouse IgG Alexa Fluor 488 or 568 (1:2000, Invitrogen); anti-mouse IgG Alexa 350 (1:2000, Invitrogen). For immunoperoxidase immunostaining, endogenous peroxidase was quenched with pre-incubation with PBS containing 3% H_2_O_2_ for 15 min. Sections were then incubated with the ABC system (Vectastain Elite ABC HRP Kit, Vector Laboratories, #PK6100). Horseradish peroxidase conjugates and 3,3′-diaminobenzidine were used for visualization of immune complexes according to the manufacturer’s manual (SeraCare, #5510-0031). Images were obtained with an Olympus BX61 microscope and analyzed with Fiji software (ImageJ). For Thioflavin-S (ThioS) labeling, sections were first mounted on slides and rehydrated prior to incubation with freshly prepared 1% Thioflavin-S (Sigma-Aldrich, #T1892) for 10 minutes at room temperature, under standard photoprotection conditions. Sections were washed 2×3 min with 80% ethanol, then 2×3 min with 95% ethanol. The final washes were performed in water 3×5 min per wash, and coverslips were placed prior to visualization.

### Cell culture

Murine primary microglia were isolated from cerebral cortices, dissected from postnatal day P0-P3 WT mice (C57BL/6 J background). Briefly, tissue was homogenized in ice-cold PBS then centrifuged at 300 x g for 5 min. The pellet was resuspended in DMEM supplemented with 10% heat-inactivated FBS (Gibco), 2 mM glutamine and penicillin/streptomycin (100 U/ml and 0.1 mg/ml respectively) and cells were seeded in poly-L-lysine T75 precoated flasks. Cultures were maintained at 37°C in humidified 95/5% air/CO_2_ incubators for 2 weeks, during which time, media change was performed twice a week. After 2 weeks, cultures were agitated at 180 rpm for 30 min to detach microglial cells from the astrocytic monolayer for collection. Isolated primary microglia were seeded at 500,000 cells per well (24 wells plate) and treated one day after with 2 μg/ml of Lipopolysaccharide (LPS, Sigma, #L3024-5MG) for 1h. Post-LPS treatment, cells were washed with PBS and RNA was extracted.

### RNA *in situ* hybridization

Mice used for RNA *in situ* hybridization (RNAscope®) were anesthetized in a CO_2_ chamber and transcardially perfused with 20 ml ice-cold PBS followed by 20 ml ice-cold 4% paraformaldehyde (PFA). Brains were then post-fixed in 4% PFA for 24h at 4ºC followed by equilibration in several sucrose gradients (10%, then 20%, finally 30%). Tissue samples were embedded in optimal cutting temperature compound (OCT), frozen on dry ice, and 10 µm-thick sections were cut using a cryostat (Leica). Fluorescent *in situ* hybridization (FISH) was performed using RNAscope^®^ according to the manufacturer’s instructions (ACD). Briefly, the mounted frozen sections were washed in PBS then baked at 60ºC for 30 min (Lab-Line Instruments Inc., Melrose Park, IL, USA). Slides were then postfixed in 4 % PFA for 1h at room temperature (RT). After fixation, the sections were dehydrated in ethanol (incubated serially in 50%, then 70%, then 100% ethanol for 5 min at each concentration) and permitted to dry at RT for 45 min. Sections were treated with H_2_O_2_ for 10 min and washed twice with distilled water. Subsequently, target retrieval was performed by boiling the slides for 5 min in retrieval reagent. Slides were then washed in distilled water, immersed in 100% ethanol, and finally subjected to air-drying for 5 min at RT. Sections were then treated with protease III for 20 min at 40ºC in the pre-warmed ACD HybEZ II Hybridization System (Cat. No. 321721, ACD) inside the HybEZ Humidity Control Tray (Cat. No. 310012, ACD). After this step, sections were washed twice with distilled water. The following probes from ACD were used and diluted at 1:50 in the RNAscope® Probe Diluent (Cat No. 300041): Tyrobp-C3 (NM_011662.2, bp8-580, Cat. No. 408191-C3), Trem2-C2 (NM_031254.3, bp2-1081, Cat. No. 404111-C2) and Apoe-C4 (NM_009696.3, bp83-1245, Cat. No. 313271-C4). Sections were then hybridized with one corresponding probe at 40ºC for 2 h in the HybEZ Oven (ACD), washed twice with 1× wash buffer, and stored overnight at RT in 5x SSC buffer (Thermo Fisher Scientific, Fair Lawn, NJ, USA). The next day, slides were rinsed 2×2 min with wash buffer (2 min per wash), followed by the three amplification steps (AMP 1, AMP 2, and AMP 3 at 40ºC for 30, 30, and 15 min respectively, with two washes with wash buffer after each amplification step). The signal was developed by treating the sections with the HRP reagent corresponding to each probe channel (e.g. HRP-C2, HRP-C3 or HRP-C4) at 40ºC for 15 min, followed by the TSA Plus fluorophore Opal 690 (dilution of 1:750 in RNAscope® Multiplex TSA Buffer [Cat. No 322809]), at 40ºC for 30 min, and HRP blocker at 40ºC for 15 min, with two wash steps after each of the incubation steps. Finally, the slides were counterstained or not with DAPI for 30 seconds. Fluorescent immunohistochemistry was performed by incubating the slides with the primary antibody diluted in PBS for 2h at RT. Slides were washed twice for 5 min per wash with 1X wash buffer, and then incubated with the secondary antibody diluted in PBS for 1h30. Immunostained slides were then washed twice with 1X wash buffer and mounted.

### Western blotting

Equal amounts of protein from (30 μg, mouse cortex was used) were separated by electrophoresis in precast 4-12% Bis-Tris Gels (Bio-Rad) and transferred to nitrocellulose membranes. The membranes were hybridized with the following primary antibodies as indicated: TYROBP (1:500, antibody provided in collaboration with Dr Richard W. Cho at Cell Signaling Technology); AT8 anti-pTAU pSer202/Thr205 (1:1000, Thermo Scientific, #MN1020); anti-GAPDH (1:2000, Santa Cruz, #SC-32233), anti-p-TAU Thr205 (1:1000, Invitrogen, #44-738G), anti-TAU HT7 (1:1000, Invitrogen, #MN1000). Secondary antibodies included: peroxidase-labeled anti-mouse IgG (1:2000, Vector Laboratories); peroxidase-labeled anti-rabbit IgG (1:2000, Vector Laboratories); peroxidase-labeled anti-rat IgG (1:2000, Vector Laboratories). ECL (Pierce, #32106) was used to reveal the immunoreactive proteins, and images were acquired using a Fujifilm ImageReader LAS-4000. Membranes were stripped using a stripping buffer (Thermo Scientific, #46430) when required. Luminescent immunoreactive protein bands were quantified using Fiji software (ImageJ).

### Aβ assays

To quantify Aβ levels, human/rat Aβ 1–40/1–42 ELISA kits (Wako, #294-64701, #290-6260) were used according to the manufacturer’s instructions. Absolute concentrations of Aβ were normalized to initial tissue weight. Supernatants from the RIPA-extracted cortices were used.

### RNA extraction and RT-qPCR analysis

RNAs were isolated from mice hippocampi using the QIAzol Lysis Reagent (Qiagen) and the miRNeasy Micro Kit (Qiagen) following the manufacturer instructions. For the qPCR analyses, 1 μg of RNA was reversed transcribed using the High Capacity RNA-to-cDNA kit (ThermoFisher, #4387406). The All-in-One qPCR Mix (GeneCopoeia, #QP001-01) was used to perform RT-qPCR. 40 cycles were done and the abundance of each transcript was normalized to the abundance of GAPDH with the ∆∆Ct method. The sequences of oligonucleotides used can be found in Litvinchuk et al. 2018^19^ except for mouse *IL6* and *TGFβ* where TaqMan® probes were used.

### RNA sequencing

RNA sequencing on mice hippocampi was performed by Novogene (https://en.novogene.com) using Illumina Novaseq 6000 S4 flow cells. Only samples with RNA integrity number (RIN) > 9 were included. Non-directional libraries were constructed with a NEB kit using the manufacturer’s protocol. RNA sequencing assays were performed after ribosomal RNA depletion by Ribo-Zero. For the data QC, four main steps were implemented including determination of the (1) distribution of sequencing quality; (2) distribution of sequencing error rate; (3) distribution of A/T/G/C bases; and (4) results of raw data filtering. The filtering process included: (1) removal of reads containing adapters, (2) removal of reads containing N > 10% (N represents bases that cannot be determined), and (3) removal of reads containing low quality bases (Qscore ≤5) that are over 50% of the total bases contained in the read. Sequencing reads were aligned to mouse reference genome mm10 (GRCm38.90) using STAR aligner guided by mouse GENCODE gene model release v15. Accepted mapped reads were summarized to gene levels using the featureCounts program. Raw count data were normalized by the voom function in the R limma package, after which differential expression was identified by the moderated t-test implemented in limma. Differentially expressed genes (DEGs) were defined to have at least 1.2-fold change in expression and Benjamini– Hochberg adjusted p ≤ 0.1 comparing different genotypes.

### Statistics

The non-genomic data were analyzed with GraphPad Prism 8. Graphs represent the mean of all samples in each group ± SEM. Sample sizes (n values) and statistical tests are indicated in the figure legends. One-way or two-way ANOVA were used for multiple comparisons. A Student’s t-test was used for simple comparisons. Significance is reported at *p < 0.05, **p < 0.01, ***p < 0.001 and ****p < 0.0001. Grubb’s test for outliers was used with α = 0.05.

## RESULTS

### *Tyrobp* transcription is increased in recruited microglia

Microglia continuously sense changes in the brain environment and are recruited to sites of injury, microbial invasion, or where abnormal folding or modification of cellular constituents are detected, as with the accumulation of aggregated Aβ. We performed dual RNA *in situ* hybridization (RNAscope^®^) and immunohistochemistry for *Tyrobp* and IBA1 respectively in WT mice and observed increased levels of *Tyrobp* mRNA in areas exhibiting clustered microglia (Fig 1A). Using the same experimental approach in two independent mouse models of cerebral amyloidosis (*APP/PSEN1*^20^ and *5xFAD*^21^), we observed a similar pattern in that the *Tyrobp* mRNA level is extensively and selectively increased in microglia recruited in close proximity to amyloid plaques as compared to microglia that are more distant from the plaques (Fig. 1B-C). We similarly assayed *Tyrobp* mRNA and IBA1 protein in the *MAPT^P301S^* mice^22^ (also known as PS19), a mutant tauopathy mouse model. We previously described an elevated number of anti-phosphorylated-TAU immunostained neurons in the piriform cortex^23^, and we detected increased amounts of *Tyrobp* mRNA in microglia in this same region (Fig. 1D). We confirmed the increase of TYROBP at the protein level in microglia around amyloid plaques (Fig. 1E) as previously reported^24^. To discriminate between the role of TYROBP in activated vs recruited microglia, we isolated primary microglia from WT mice and exposed them to the gram-negative bacterial endotoxin lipopolysaccharide (LPS) to induce microglial activation^25^, the status of which we established by quantifying the robust increase of *Tnfα* mRNA following LPS treatment. Interestingly, *Tyrobp* mRNA level was unchanged, suggesting that *Tyrobp* transcription may be increased only when microglia are *both* recruited *and* activated but not in resident microglia despite evidence that these residents are also activated (Fig. 1F).

**Figure 1:**
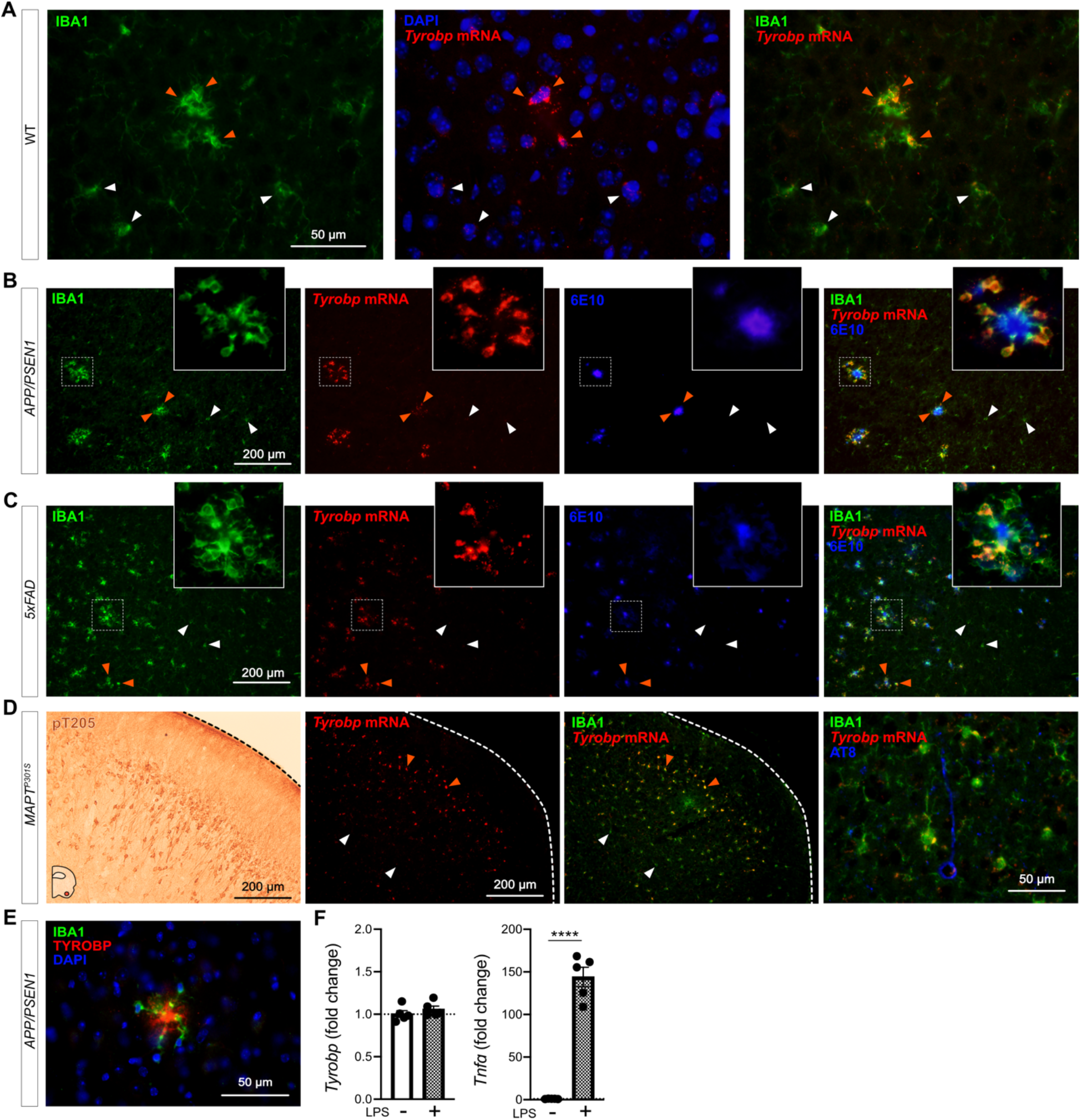
*Tyrobp* mRNA is increased in recruited microglia. (A) Dual RNA fluorescent *in situ* hybridization (RNAscope®) and immunohistochemistry for *Tyrobp* mRNA (red) and IBA1 protein (green) respectively in WT mice (DAPI in blue). Scale bar = 50 μm. (B-C) Dual RNA *in situ* hybridization and immunohistochemistry for *Tyrobp* (red), IBA1 (Green) and Aβ (antibody 6E10) (blue) in *APP/PSEN1* (B) and *5xFAD* (C) mice. Scale bar = 200 μm. (D) Left panel: representative image of immunohistochemistry with antibody pT205 in the piriform cortex of *MAPT^P301S^* (PS19) mice. Scale bar = 200 μm. Right panels: dual RNA *in situ* hybridization and immunohistochemistry for *Tyrobp* (red), IBA1 (green) and pTau (antibody AT8) (blue) in the piriform cortex of *MAPT^P301S^* mice. Scale bars = 200 and 50 μm. (E) Co-immunohistochemistry for TYROBP (green) and human Aβ (antibody 6E10) (red) in *APP/PSEN1* mice (DAPI in blue). Scale bar = 50μm. (F) RT-qPCR analyses of *Tyrobp* and *TNFα* mRNAs in WT primary microglia with and without LPS. Mice were either 4 (A) or 8 (B-E) months of age and were all WT for *Tyrobp*. White and orange arrows indicate examples of non-recruited and recruited microglia respectively. Slice thickness = 10μm.

### Microglia are normal in *Iba1*^*Tyrobp*^ mice

To determine whether constitutive elevation of TYROBP via transgenesis may influence microglial phenotype and progression of AD pathology, we generated a novel transgenic mouse overexpressing *Tyrobp* selectively in microglia in the central nervous system. We used the mouse *Tyrobp* and *Enhanced Green Fluorescent Protein* (EGFP) sequences separated by an Internal Ribosome Entry Site (IRES) under the control of the mouse *Iba1* regulatory sequences (Supp Fig. 1^17^). Microinjections were performed in C57BL/6J mice and one line was selected for further use based on expression level of the transgene, now referred to *Iba1^Tyrobp^*. We first assessed the overexpression of *Tyrobp* mRNA by RT-qPCR and measured a ≈ 2.5-fold increase (Fig. 2A). Despite this elevated mRNA level, western blot analyses of protein extracts from the cortex did not reveal a significant overexpression at the protein level (Fig. 2B). Using combined RNA *in situ* hybridization and immunohistochemistry for *Tyrobp* and IBA1, respectively, in mice deficient (*Tyrobp^−/−^*), WT or overexpressing (*Iba1^Tyrobp^*) *Tyrobp*, we confirmed the 2-fold increase in *Tyrobp* mRNA in *Iba1^Tyrobp^* mice compared to WT (Fig. 2C-D). We observed that only a subset of microglia was overexpressing *Tyrobp* mRNA in *Iba1^Tyrobp^* mice. This selectivity is likely due to the use of the *Iba1* promoter in a WT background without extensive microglia activation, thereby also accounting for the lack of a significant increase of TYROBP at the protein level in the resting state. RNA sequencing of hippocampi from *Iba1^Tyrobp^* mice did not reveal any differentially expressed genes (DEGs) other than *Aif1* (= *Iba1*), which is increased due to the inclusion of the first two exons in the transgenic vector (Supp Fig. 1). These data indicate that *Iba1^Tyrobp^* microglia do not display molecular and phenotypic changes in WT mice.

**Figure 2:**
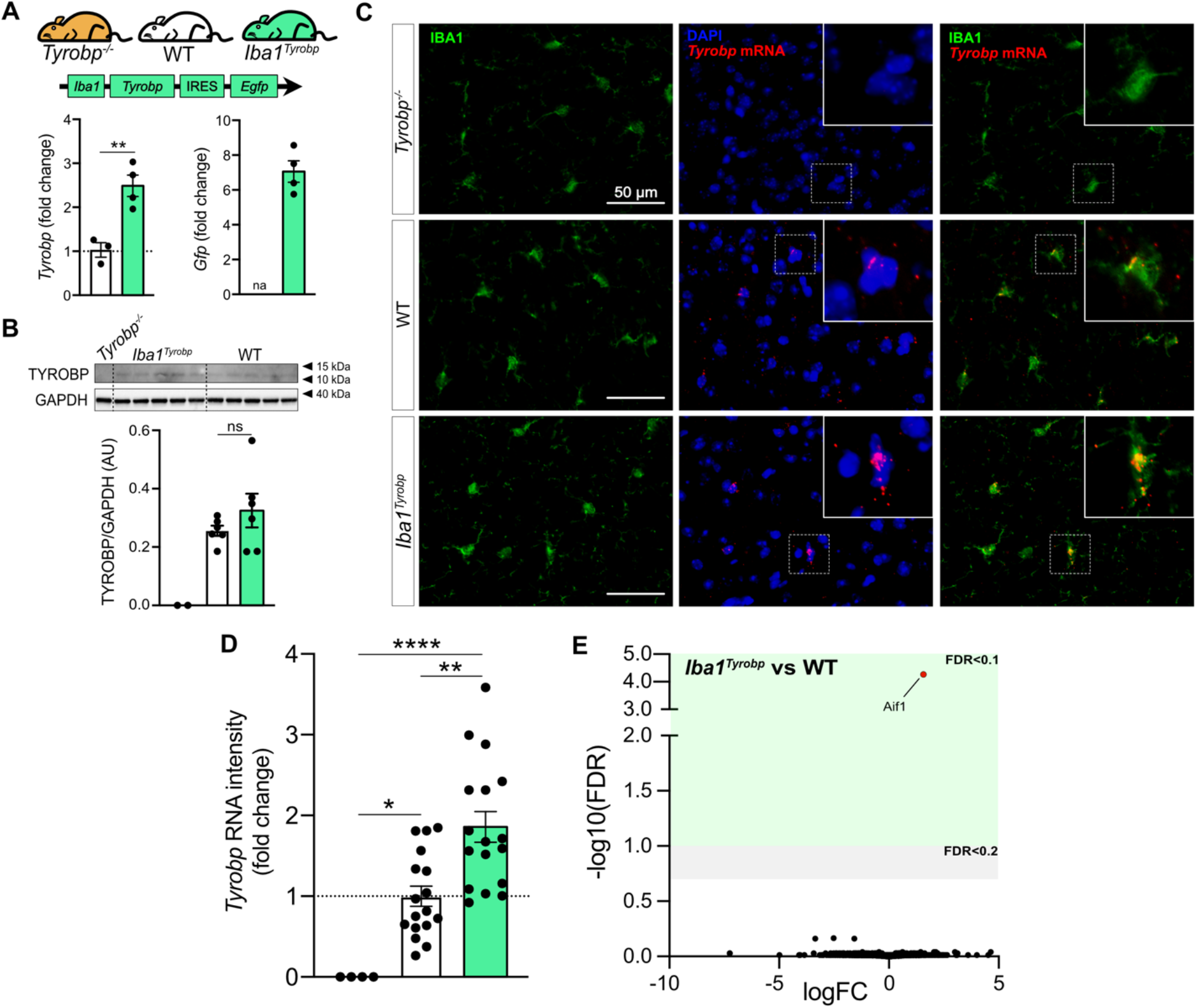
Generation of *Iba1^Tyrobp^* mice. (A) Hippocampi from 4-month old *Tyrobp^−/−^*, WT and *Iba1^Tyrobp^* mice were assayed for *Tyrobp* and *Gfp* mRNAs by RT-qPCR (n = 3-4 mice per group). (B) Representative western blot and quantification of TYROBP and GAPDH in the cortex of the same groups used in (A) (n = 2-6 mice per group). (C) Dual RNA fluorescent *in situ* hybridization and immunohistochemistry for *Tyrobp* mRNA (red) and IBA1 (green) respectively (DAPI in blue) in Tyrobp^−/−^, WT and *Iba1^Tyrobp^* mice. Scale bar = 50 μm and slice thickness = 10μm. (D) Quantification of *Tyrobp* mRNA intensity from the experiment described in (C). n = 4, 17 and 17 slices per group (from N = 1 mouse per genotype) for *Tyrobp^−/−^*, WT and *Iba1^Tyrobp^* mice respectively. (E) Volcano plot representation of the whole hippocampal DEGs in *Iba1^Tyrobp^* vs WT mice (n = 4 4-month old males per genotype). Error bars represent means ± SEM. Statistical analyses were performed using a Student t-test (A) or a One-Way ANOVA followed by a Tukey’s post-hoc test (B, D), *p < 0.05, **p < 0.01, ****p < 0.0001. na = not applicable; ns = non-significant.

### TYROBP overexpression in microglia decreases amyloid plaque load in *APP/PSEN1* mice

To assess whether TYROBP overexpression in microglia modulates Aβ deposition in *APP/PSEN1* mice, double-heterozygous *APP/PSEN1;Iba1^Tyrobp^* mice were generated and studied at 4 months of age. We measured a ≈50% decrease of the plaque density in the cerebral cortices of both male and female *APP/PSEN1;Iba1^Tyrobp^* mice compared to sex-matched *APP/PSEN1* mice (Fig. 3A-B) in sections stained for amyloid plaques using Thioflavin S (ThioS). This observation was supported by measuring levels of human Aβ42 and Aβ40 by ELISA, both of which were apparently associated with a trend toward decreases in the cortex with TYROBP overexpression, mostly among male *APP/PSEN1;Iba1^Tyrobp^* mice (Fig. 3C). There was no genotype-dependent difference in the number of plaque-associated microglia (Fig. 3D-E), unlike what has been reported in *5xFAD* mice in the presence of a transgenic increase in TREM2^26^. To evaluate microglia activation, we probed both groups with anti-IBA1 antibody and observed a weaker staining in *APP/PSEN1;Iba1^Tyrobp^* mice (Supp Fig. 2A). We next performed RT-qPCR on a group of microglial genes previously described in homeostatic or activated microglia. There was a significant increase of *Axl* and *Ccl2* and a decrease of *Cd68* and *Tgfβ* in brains of *APP/PSEN1;Iba1^Tyrobp^* mice (Fig. 3F). Finally, we observed that the decrease of amyloid plaques persisted in 8-month-old *APP/PSEN1;Iba1^Tyrobp^* mice as shown by the decreased percentage of ThioS positive areas in somatomotor, hippocampus, and visual areas (Fig. 3G-H).

**Figure 3:**
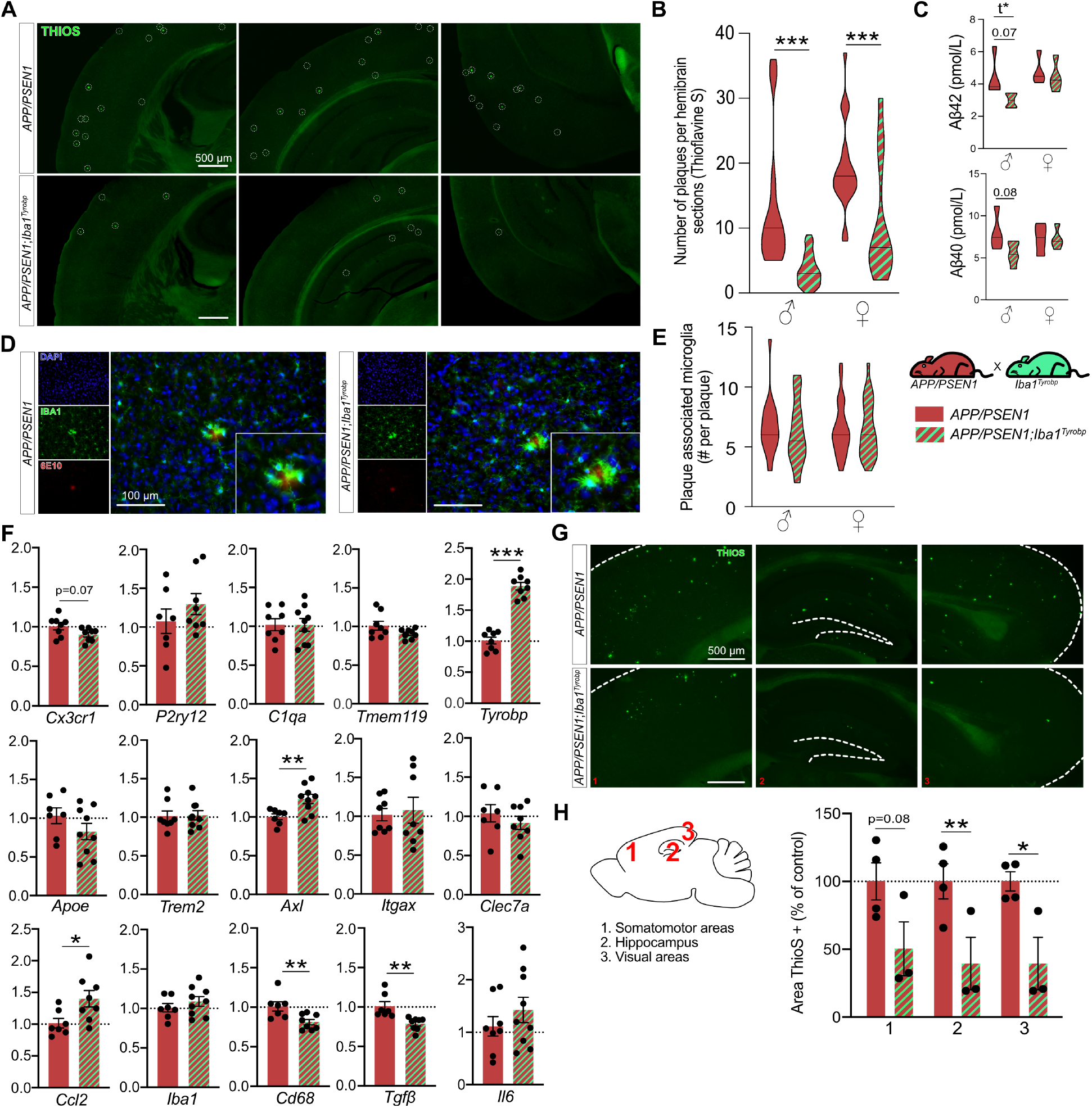
Transgene-derived *Tyrobp* upregulation decreases amyloid plaque load in *APP/PSEN1* mice. (A) Representative images of Thioflavine-S (ThioS) staining in *APP/PSEN1* and *APP/PSEN1;Iba1^Tyrobp^* mice at 4 months of age. Scale bar = 500 μm. (B) Quantification of the number of ThioS-positive plaques per hemibrain in *APP/PSEN1* and *APP/PSEN1;Iba1^Tyrobp^* mice at 4 months of age. N = 4-5 mice per genotype and sex with 3 slices per animal. (C) Human Aβ42 and Aβ40 concentrations measured by ELISA in the cortex of the same groups described in (B). (D) Representative images of double-label immunohistochemistry with anti-IBA1 and anti-6E10 antibodies in *APP/PSEN1* and *APP/PSEN1;Iba1^Tyrobp^* mice at 4 months of age. Scale bar = 100 μm. (E) Quantification of the number of plaque-associated microglia in the 4 groups described in (B). N = 10-24 plaques from 4-5 mice per group. (F) RT-qPCR analyses of microglial gene mRNAs in the hippocampus of *APP/PSEN1* and *APP/PSEN1;Iba1^Tyrobp^* mice at 4 months of age. N = 7-9 mice per group, females and males were pooled. (G) Representative images of ThioS staining in male *APP/PSEN1* and *APP/PSEN1;Iba1^Tyrobp^* mice at 8 months of age. Scale bar = 500 μm. (H) Quantification of the ThioS immunoreactive area in male *APP/PSEN1* and *APP/PSEN1;Iba1^Tyrobp^* mice at 8 months of age (somatomotor and visual areas of the cortex, and hippocampus were quantified). N = 3-4 mice per group. Error bars represent means ± SEM. Statistical analyses were performed using a Two-Way ANOVA followed by a Sidak post-hoc test for (B,C and E) or a Student t-test for (C) when *t is indicated and (F-H), *p<0.05, **p<0.01, ***p<0.001.

### TYROBP overexpression in *MAPT*^*P301S*^ mice increases TAU phosphorylation and microglia activation

We previously reported that deletion of *Tyrobp* altered both mouse amyloid and tauopathy phenotypes and the microglial response to these pathologies^23, 27, 28^. In *MAPT^P301S^;Iba1^Tyrobp^* double heterozygous mice, western blot analyses using AT8 and T205 antibodies revealed increased levels of phosphorylated-TAU (pTAU) in the cortex of both male and female mice compared to *MAPT^P301S^* mice at 4 months of age, whereas total human TAU levels detected with the HT7 antibody were unchanged (Fig. 4A-B). Increased pTAU within brains from *MAPT^P301S^*;*Iba1^Tyrobp^* mice was further confirmed immunohistochemically (Fig. 4C). We also observed increased IBA1 intensity in *MAPT^P301S^*;*Iba1^Tyrobp^* compared to *MAPT^P301S^* mice that correlated with the increased pTAU (Fig. 4D, Supp Fig. 2B). We confirmed an increased microglial activation state by double-label immunohistochemistry with anti-IBA1 and anti-CD68 in the piriform cortex (Fig. 4E). Using RT-qPCR, we measured increases of *Tyrobp, P2ry12*, *Apoe*, *Axl*, *Itgax*, *Iba1*, *Tgfβ* and *Il6 mRNAs* in *MAPT^P301S^*;*Iba1^Tyrobp^* mice compared to *MAPT^P301S^* mice (Fig. 4F).

**Figure 4:**
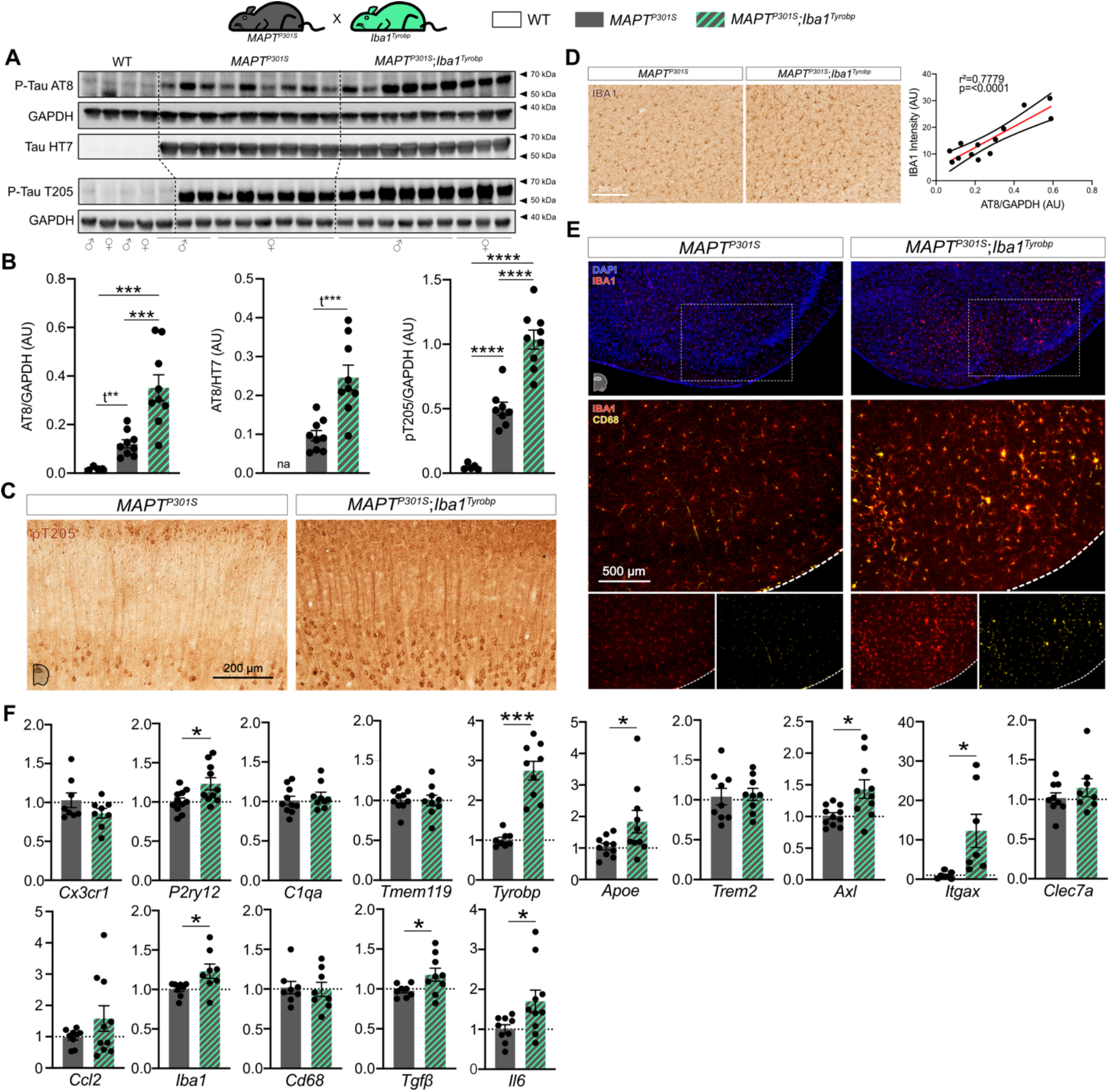
Transgene-induced *Tyrobp* upregulation increases TAU hyperphosphorylation and microglial activation in 4-month-old *MAPT^P301S^* mice. (A) Western blot analyses of phosphorylated-TAU on S202 or T205 epitopes (AT8 and pT205 antibodies) and total human TAU (HT7 antibody) in cortical homogenates from WT, *MAPT^P301S^* (PS19) and *MAPT^P301S^;Iba1^Tyrobp^* mice at 4 months-old. n = 4-9 mice per group. (B) Densitometric analyses of western blots presented in (A) standardized to GAPDH or HT7. (C) Representative images of DAB-immunohistochemistry with antibody pT205 in 4 month-old *MAPT^P301S^* and *MAPT^P301S^;Iba1^Tyrobp^* mice. Scale bar = 200 μm. (D) Left panel: representative images of anti-IBA1 immunohistochemistry on the same groups described in (C). Scale bar = 200 μm. Additional representative pictures are presented in Supplementary Figure 2. Right panel: western blot-AT8/GAPDH quantification plotted against anti-IBA1 immunoreactivity in the cortex. Linear regression with trend line (red line) and 95% confidence intervals (black lines) are indicated. (E) Representative images of double-label immunofluorescence with anti-IBA1 and anti-CD68 antibodies in the piriform cortex on the same groups described in (C). Scale bar = 500 μm. (F) RT-qPCR analyses of microglial gene mRNAs in the hippocampus of *MAPT^P301S^* and *MAPT^P301S^;Iba1^Tyrobp^* mice at 4 months of age. N = 7-11 per group. Error bars represent means ± SEM. Statistical analyses were performed using a One-Way ANOVA followed by a Tukey’s post-hoc test for (B) or a Student t-test for (B) when *t is indicated and (F), *p<0.05, **p<0.01, ***p<0.001, ****p<0.0001.

### *Iba1*^*Tyrobp*^ mice reveal a likely relationship between *Tyrobp* and *Apoe*

Despite the obvious differences across *APP/PSEN1* and *MAPT^P301S^* mouse models and the diverse consequences of *Tyrobp* upregulation in each of these mice, there are shared changes in *Axl*, *Ccl2*, *Tgfβ* and *Il6* mRNAs in both *APP/PSEN1* or *MAPT^P301S^* mice overexpressing TYROBP (Fig. 3F and 4F). These genes have been recently associated with *Apoe* in microglia, macrophages and mononuclear phagocytes. AXL has been identified as a regulator of APOE^29^ and accumulation of IL6 and CCL2 have been associated with APOE overexpression^30,31^. Similarly, reciprocal suppression of TGFβ and induction of APOE have been described in DAM microglia^15^. *Apoe* is indeed significantly upregulated in *MAPT^P301S^*;*Iba1^Tyrobp^* mice as compared to *MAPT^P301S^* mice but not in *APP/PSEN1;Iba1^Tyrobp^* mice compared to *APP/PSEN1* mice (Fig. 3F and 4F). However, in bulk RNA sequencing on hippocampi from male *APP/PSEN1;Iba1^Tyrobp^* vs *APP/PSEN1* mice, we identified *Apoe* as a potential (activation z-score: 2.44; p-value overlap: 0.224) upstream regulator (Supp Fig. 3), suggesting the possible existence of a relationship between *Tyrobp* upregulation and *Apoe*. While a TREM2-APOE pathway has been described^15^, it is interesting to note that these data support the possibility that the TYROBP-APOE relationship is detectable even in the absence of *Trem2* upregulation (Fig. 3F and 4F).

### Induction of microglial *Tyrobp* and *Apoe* is *Trem2*-independent in a model of cortical stab injury

To assess the interactions amongst *Trem2*, *Tyrobp* and *Apoe* in microglia, we used an injury paradigm by introducing a small penetrating cortical stab injury via stereotactic surgery into one hemisphere of the mouse brain in order to induce a recruitment of microglia around the injury site^32^. We first used injured-WT mice and combined RNA *in situ* hybridization and immunohistochemistry for *Apoe* and IBA1 respectively. In the intact hemisphere, most *Apoe* mRNA was not located in microglia but rather in astrocytes, the source of most APOE in brain. However, *Apoe* mRNA was dramatically increased in microglia recruited on the lesioned side (Fig. 5A). Following the same procedure in *Tyrobp^−/−^* mice, *Apoe* mRNA was not induced in microglia on either side (Fig. 5B), but strikingly, mRNA levels of *Tyrobp* and *Apoe* were highly upregulated in the recruited microglia of injured *Trem2^−/−^* mice (Fig. 5C, Supp Fig. 4). Taken together, these data indicate that *Tyrobp* upregulation in recruited microglia around the traumatic lesion is *Trem2*-independent. Moreover, the increase of *Apoe* transcripts in recruited microglia in the same mouse model of injury appears to be *Tyrobp-*dependent and *Trem2-*independent.

**Figure 5:**
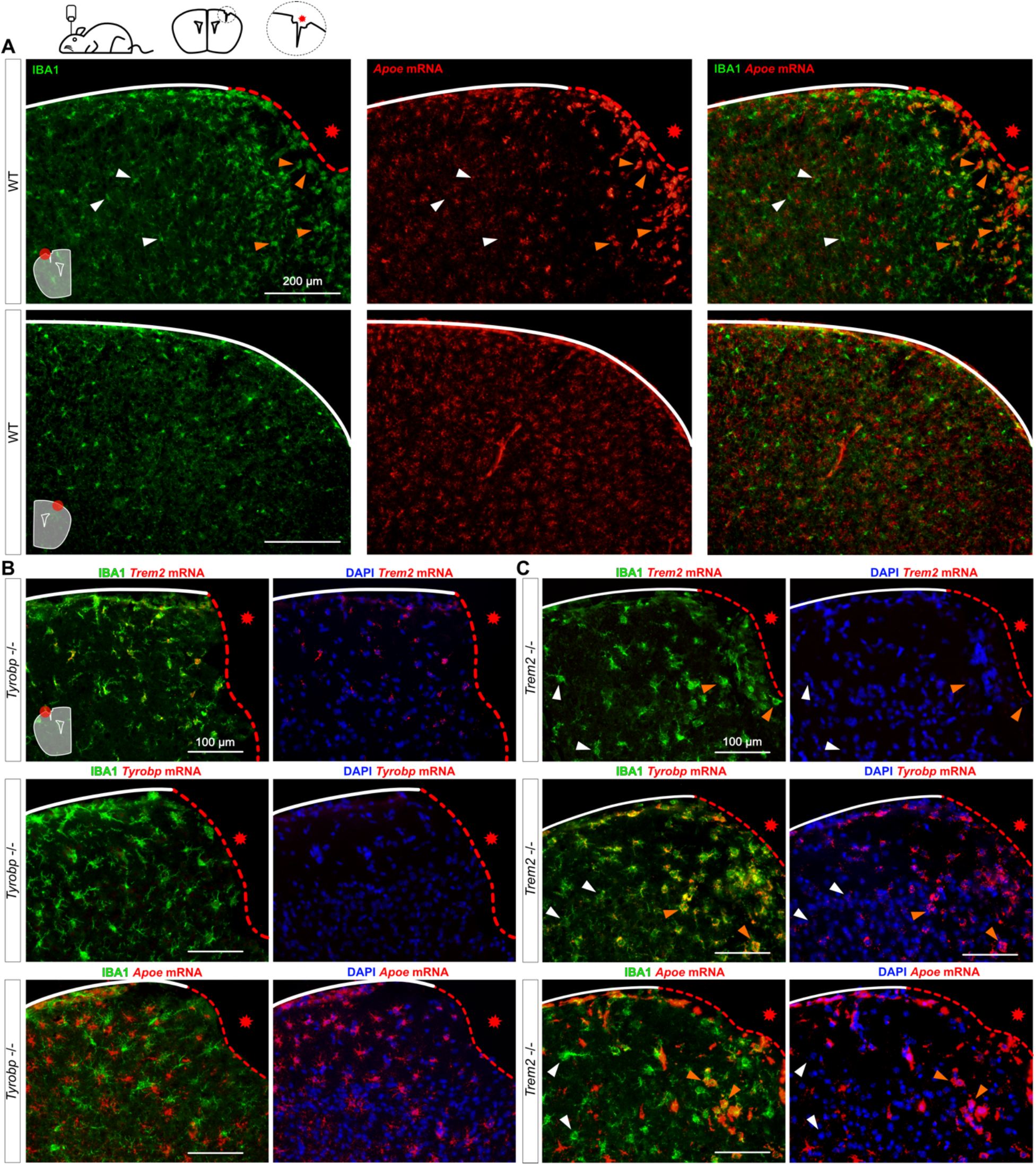
Increases of *Tyrobp* and *Apoe* mRNAs in microglia recruited toward a site of stab injury are *Trem2*-independent. (A) Stab-injured WT mice were sacrificed 3 days after injury and dual RNA fluorescent *in situ* hybridization and immunohistochemistry for *Apoe* mRNA (red) and anti-IBA1 (green) respectively was performed. The injured ipsilateral area (red dotted line) is shown on the top row and the uninjured contralateral area is show on the bottom row. Scale bar = 200 μm. (B-C) The same stab injury protocol was utilized in *Tyrobp^−/−^* (B) and *Trem2^−/−^* (C) mice. Anti-IBA1 and DAPI stainings are shown in green and blue, respectively. Top row: *Trem2* mRNA (red); middle row: *Tyrobp* mRNA (red); bottom row: *Apoe* mRNA (red). Mice were 4 months of age and slice thickness = 10 μm. The red asterisk indicates the injured side. White and orange arrows indicate examples of non-recruited and recruited microglia, respectively.

### Induction of microglial *Tyrobp* and *Apoe* around amyloid plaques is *Trem2*-independent, and *Apoe* upregulation is dramatically decreased when *Tyrobp* is absent

In order to investigate further these interactions among *Trem2*, *Tyrobp* and *Apoe* in microglia in the presence of mutant human *APP*, we first performed dual RNA *in situ* hybridization and immunohistochemistry for *Tyrobp*, IBA1 and 6E10 in *TgCRND8* mice^33^ on either a WT or *Trem2*-null background. Despite reduced recruitment of microglia around the plaques when *Trem2* is deleted^24^, *Tyrobp* mRNA was still increased in plaque-associated microglia (Fig. 6A). as was *Apoe* mRNA in plaque-associated microglia in the same *TgCRND8*;*Trem2*^−/−^ mice (Fig. 6B). We then assayed *APP/PSEN1* mice that were either WT or deficient for *Tyrobp* and, while the expression of *Apoe* was not completely abolished by deletion of *Tyrobp*, we confirmed a substantial decrease in the induction of *Apoe* mRNA in plaque-associated microglia when *Tyrobp* was absent (Fig. 6C).

**Figure 6:**
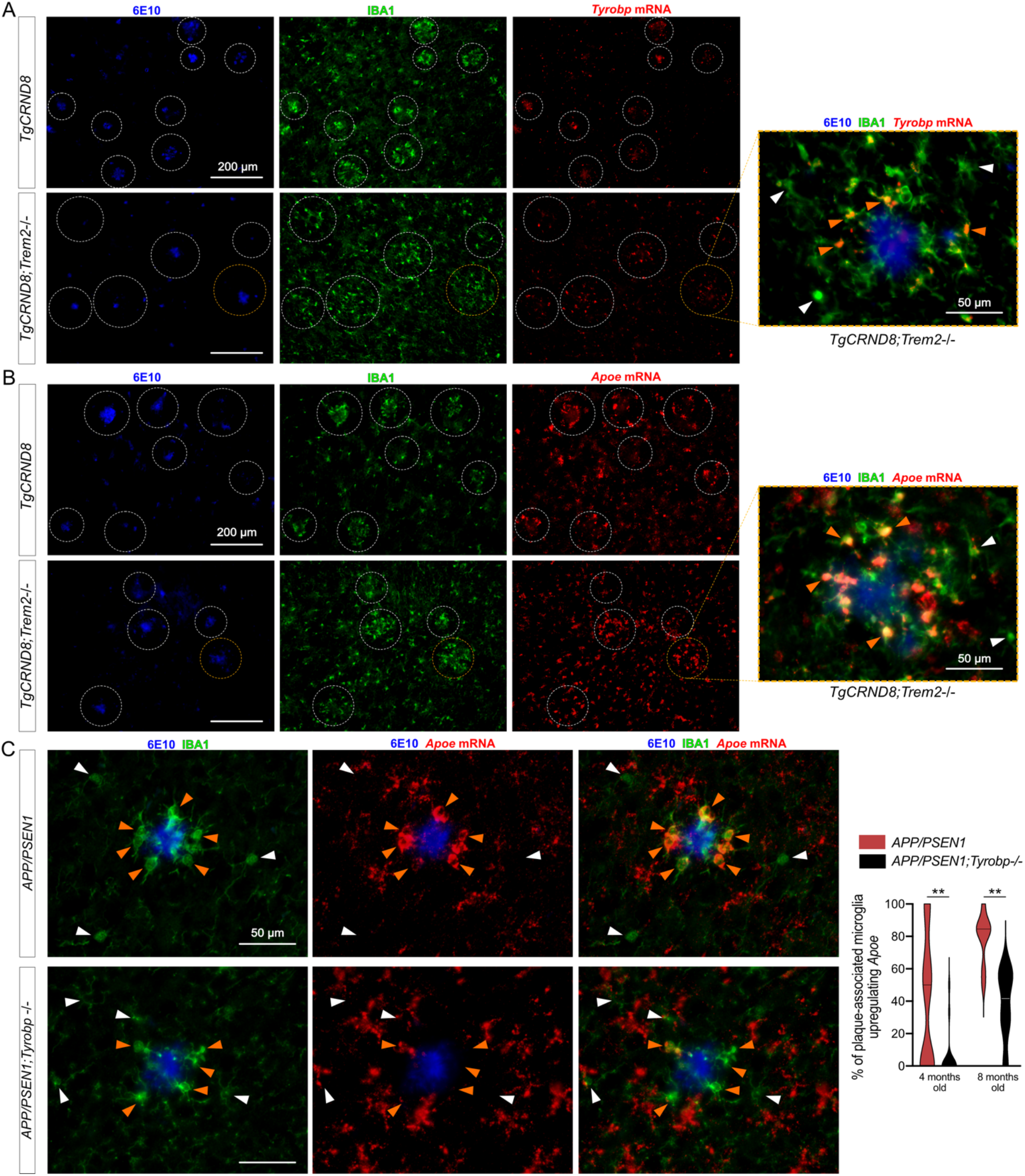
Increases in *Tyrobp* and *Apoe* mRNAs in amyloid plaque-associated microglia are *Trem2* independent. (A) Dual RNA fluorescent *in situ* hybridization and immunohistochemistry for *Tyrobp* mRNA (red), anti-IBA1 (green) and human Aβ (6E10 antibody) (blue) in *TgCRND8* mice on WT (top row) or *Trem2*^−/−^ (bottom row) background. Scale bar = 200 or 50 μm. (B) Dual RNA fluorescent *in situ* hybridization and immunohistochemistry for *Apoe* mRNA (red), anti-IBA1 (green) and human amyloid (6E10 antibody) (blue) in the same mice as in (A). Scale bar = 200 or 50 μm. (C) Dual RNA fluorescent *in situ* hybridization and immunohistochemistry for *Apoe* mRNA (red), anti-IBA1 (green) and human Aβ (6E10 antibody) (blue) in *APP/PSEN1* mice on a WT (top row) or *Tyrobp*-null (bottom row) background. Scale bar = 50 μm. Right panel: quantification of the number of plaque-associated microglia with upregulated *Apoe* mRNA in the same mice as in (D). N = 2-3 mice per group (A-C).

## DISCUSSION

*TREM2*, *TYROBP* and *APOE* are three microglial genes linked in a pathway contributing to the pathogenesis of AD and in the transition to DAM^10^. *TYROBP* was identified as a key driver in late onset AD^9^. TREM2 binds to TYROBP, its intracellular adaptor, to initiate its signal transduction pathway, and naturally occurring loss-of-function mutations of either TYROBP or TREM2 can lead to Nasu-Hakola disease^13,34^. There is a general assumption among investigators in this research area that genetic deletion or overexpression of either TREM2 or TYROBP would result in identical phenotypes in disease models, but, until now, this has not been tested directly. We previously demonstrated amelioration of behavior and electrophysiologic impairments in *APP/PSEN1* and *MAPT^P301S^* mice on a *Tyrobp*-null background, despite a concurrent absence of effect on amyloid pathology and an apparent increase in the stoichiometry of phosphorylated TAU vs total TAU^23, 27, 28^. Homozygous deletion of *Trem2* can also lead to amelioration of both amyloidosis and tauopathy ^35,36^, but those effects vary according to the mouse model, and the age and level of deficiency at sacrifice ^23, 37, 38^. Lee *et al.*^26^ used bacterial artificial chromosome (BAC)-mediated transgenesis to overexpress the human *TREM2* in the mouse genome and showed that TREM2 overexpression reduces amyloid accumulation in 5xFAD mice. Using a similar BAC system, Gratuze *et al.*^39^ assessed the impact of *TREM2^R47H^* in *MAPT^P301S^* mice but no TREM2 overexpression was reported in that study.

The possible existence of an early TREM2-independent phase in conversion of microglia to DAM was described by Keren-Shaul *et al.^10^* but was not evident in studies by either Krasemann *et al.*^15^ or Zhou *et al.*^40^ *Apoe* has also been described as a participant in stage 1 of DAM with *Tyrobp*^10^ and it has been suggested that it drives the DAM transition through a TREM2-APOE pathway^15^. Moreover, *Apoe* has been reported to influence both amyloidosis and tauopathy histological phenotypes in mouse models^41,42^. A complete elucidation of the choreography of the regulatory interactions among these genes and their cognate proteins therefore remains an area of intense interest. We would suggest that the discrepancies across the various analyses might be explained in part by the fact that DAM microglia are located in the immediate proximity of the plaques, and that neither bulk-nor single-cell-RNA sequencing can distinguish homeostatic vs DAM phentoypes since both techniques generate an average transcriptome analysis from all microglia in a given tissue sample. This formulation played a major role in prompting us to use dual RNA *in situ* hybridization and immunohistochemistry in the current study in which we sought to determine 1) the effects of transgenic overexpression of *Tyrobp* on amyloid and tau pathologies and 2) the relationship of the induction of *Tyrobp* to these pathologies and to the induction of *Trem2* and *Apoe*.

While this manuscript was in preparation, Chen *et al*. (2020) reported spatial transcriptomics and *in situ* sequencing in *App^NL-G-F^* mice to avoid the averaging of the transcriptome in a tissue sample. These investigators proposed the formulation of a plaque-induced gene (PIG) network in microglia and astrocytes in immediate proximity to amyloid plaques with differential expression of 57 genes^43^. Top genes highly upregulated in the proximity of the plaques and as early as 3 months of age were *Tyrobp*, *Apoe* and several complement-related genes. Notably, *Tyrobp* is a key regulator of the complement subnetwork^9^ and C1q is down-regulated in *APP/PSEN1* and *MAPT^P301S^* mice in the absence of *Tyrobp*^23,28^. The authors performed the same type of experiment in human AD brain slices and confirmed the enrichment of *Tyrobp* and several complement components (*C1qA*, *C1qB*, *C1qC*, and *Clu*). Of particular relevance to our data herein, *Trem2* was not included in the human PIG network. Furthermore, Srinivasan *et al.*^44^ recently used RNA sequencing to profile fluorescence-activated cell sorted (FACS)-purified human microglia from frozen AD and control brains and showed that one of the top gene upregulated in human-AD microglia was *Apoe*, confirming potential importance of the interplay between *Tyrobp* and *Apoe*.

With the inclusion of *Tyrobp* as a PIG gene^43^ and because of our prior validation of its actions as a driver of AD^9,23,27,28^, we hypothesized that constitutive overexpression of microglial *Tyrobp* via transgenesis would alter both amyloid and tau pathologies. In *APP/PSEN1* mice, *Tyrobp* upregulation decreased the amyloid burden, similar to what occurs with the upregulation of *Trem2*^26^ and supported by a recent report showing that higher microglia activation shows protective effects on subsequent amyloid accumulation^45^. In *MAPT^P301S^* mice, we previously reported that deficiency of *Tyrobp* increased TAU phosphorylation and spread^23^, and were puzzled, therefore, to observe a similar increase in TAU phosphorylation in the *MAPT^P301S^* mice with overexpression of *Tyrobp*. *MAPT^P301S^*;*Iba1^Tyrobp^* microglia are more reactive than *MAPT^P301S^* microglia, and reactive microglia have been reported to drive TAU pathology, so these two observations are compatible^46,47^. This exacerbation of pathology under conditions of either down- or up-regulation of *Tyrobp* in *MAPT^P301S^* mice reveals the complexity of the microglial events underpinning tauopathy. Our formulation is that, for any particular microglial activation status, there exists some optimum level of *Tyrobp* expression, and that either elevation or deficiency of *Tyrobp* levels can be detrimental. These data also support a key role for microglial TYROBP in AD pathology progression, as we previously proposed in our reports on mice deficient for *Tyrobp*^23,27,28^.

Our data indicate that *Tyrobp* upregulation is an early marker of recruited microglia and can occur even in the brains of *Trem2-*deficient mice. Similarly, we observed that the increased *Apoe* mRNA level in microglia is *Trem2*-independent, whether in injury or AD-related mouse models. Finally, we observed that microglial *Apoe* mRNA level was greatly attenuated in plaque-associated microglia in *Tyrobp*-deficient mice. These data confirm the model proposed by Keren-Shaul *et al.*^10^ in which *Tyrobp* and *Apoe* transcripts are increased first, and neither transcription event requires the presence of *Trem2*. Moreover, Meilandt *et al.*^48^ recently reported that microglial APOE expression was not reduced, but, on the contrary, was increased in *PS2APP;Trem2^−/−^* when compared with microglial APOE expression in *PS2APP* mice. In that same study, they also analyzed the expression profiles of FACS-purified microglia from *5xFAD* mice either in the presence or absence of *Trem2* and showed a two-fold reduction in *Apoe* in one dataset (GSE132508)^10,48^ but no reduction at all in the other (GSE65067)^48,49^. However, Parhizkar *et al.*^50^ reported that the absence of functional TREM2 reduces plaque-associated APOE. This is in line with what Krasemann *et al.*^15^ proposed when they showed that genetic targeting of *Trem2* suppresses the APOE pathway. Our observations and conclusions herein apparently differ from those of Parhizkar *et al.*^50^ to the extent that, in our hands, microglial amyloid-plaque sensing followed by upregulation of *Tyrobp* and *Apoe* are preserved despite the absence of *Trem2* and, as a consequence of the *Trem2* deficiency, microglia recruitment into the proximity of amyloid plaques is reduced. This relationship points to the fact that the absence of functional TREM2 will block appearance of the full DAM phenotype and therefore the associated clearance of the plaques is reduced. Nevertheless, we propose a model wherein the sensing of amyloid plaques --which takes place upstream of amyloid plaque clearance--involves *Tyrobp* and *Apoe* but not necessarily *Trem2*.

Considering the central role of *Tyrobp* described in the microglial sensome^8^, its upregulation even in the absence of *Trem2*, and the consequences of its overexpression in *APP/PSEN1* and *MAPT^P301S^* mice, we propose that *Tyrobp* is one of the key genes upregulated in the switch from the homeostatic phenotype to the DAM phenotype. Taking all the data together, we propose a model in which microglia perceive stimuli and initiate their responses during an early and relatively brief time window, corresponding to what Keren-Shaul *et al*.^10^ described as stage 1. During this phase, *Trem2* is not required but *Apoe* will be upregulated presumably as a downstream consequence of *Tyrobp* upregulation. TYROBP is a 113 amino acid polypeptide with a minimal extracellular region^51,52^, making it unlikely that TYROBP is the sole player in a signal transduction pathway involving both the perception of the environment and the triggering of the switch from homeostatic phenotype to DAM. However, TYROBP is the adaptor for many receptors other than TREM2^16^, and therefore, it is plausible and perhaps likely that other TYROBP receptors could play key roles in sensing the deposition of amyloid. For example, numerous SIGLEC proteins (sialic acid-binding immunoglobulin-type lectins) carry a positively charged residue in their transmembrane domain that participates in oligomerization of the SIGLEC with TYROBP. The primary SIGLEC ligand is a sialic acid that accumulates in many pathological conditions including cerebral Aβ amyloidosis^53,54^. Moreover, Siglec-H interacts with TYROBP, and its expression has been reported to be elevated in *5xFAD* mice vs WT^26^. CD33 (SIGLEC-3) is also one of the most abundant SIGLECs in human brain, and genome-wide association studies (GWAS) implicated a polymorphism near *CD33* as a genetic risk factor for AD^2, 4, 55^. CD33 and TREM2 both interact with TYROBP, either directly (TREM2) or via common intracellular signaling factors (CD33). Griciuc *et al.*^56^ recently investigated crosstalk between CD33 and TREM2 and proposed that CD33 acts upstream of TREM2. They also showed that *Cd33* and *Tyrobp* expression levels did not change in *Trem2^−/−^* versus WT microglia. This formulation provides evidence that CD33-TYROBP signaling could occur upstream of the recruitment and upregulation of TREM2.

Rather than the somewhat “*Trem2*-centric” view of DAM proposed in the existing AD microglia literature^15, 40, 50^, we propose that *Tyrobp* plays a central role in an alternative and early pathway in the microglial sensome^8^, even in the absence of any change in *Trem2* levels. The data that we present here document the robust consequences of TYROBP overexpression in both *APP/PSEN1* and *MAPT^P301S^* mice. We confirm here that upregulation in microglia of both *Tyrobp* and *Apoe* constitute interconnected events in microglia sensing of amyloid deposits, and these events take place independently of *Trem2*.

## AUTHOR CONTRIBUTIONS

MA, JVHM, SG and MEE designed the study. MA performed the experiments and analyzed the data. JM contributed to the RNA *in situ* hybridization related experiments. MW and BZ contributed to the RNA sequencing analysis. JKG and PF provided the *TgCRND8* mice. MA, SG and MEE wrote the manuscript.

---

**Supplementary Figure 1:**
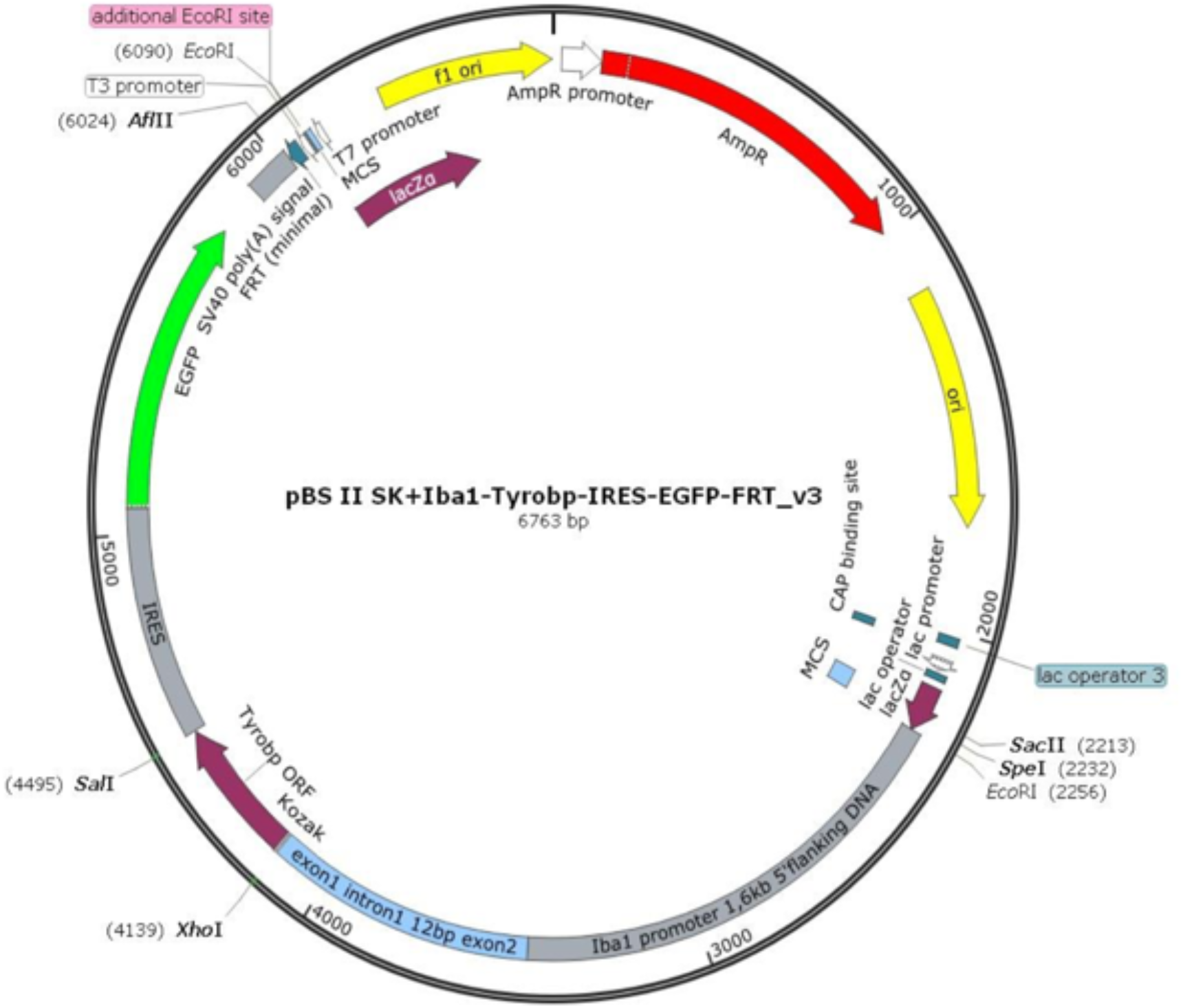
Plasmid used to generate the *Iba1^Tyrobp^* transgenic mice. The insert was linearized after digestion with *PacI*.

**Supplementary Figure 2:**
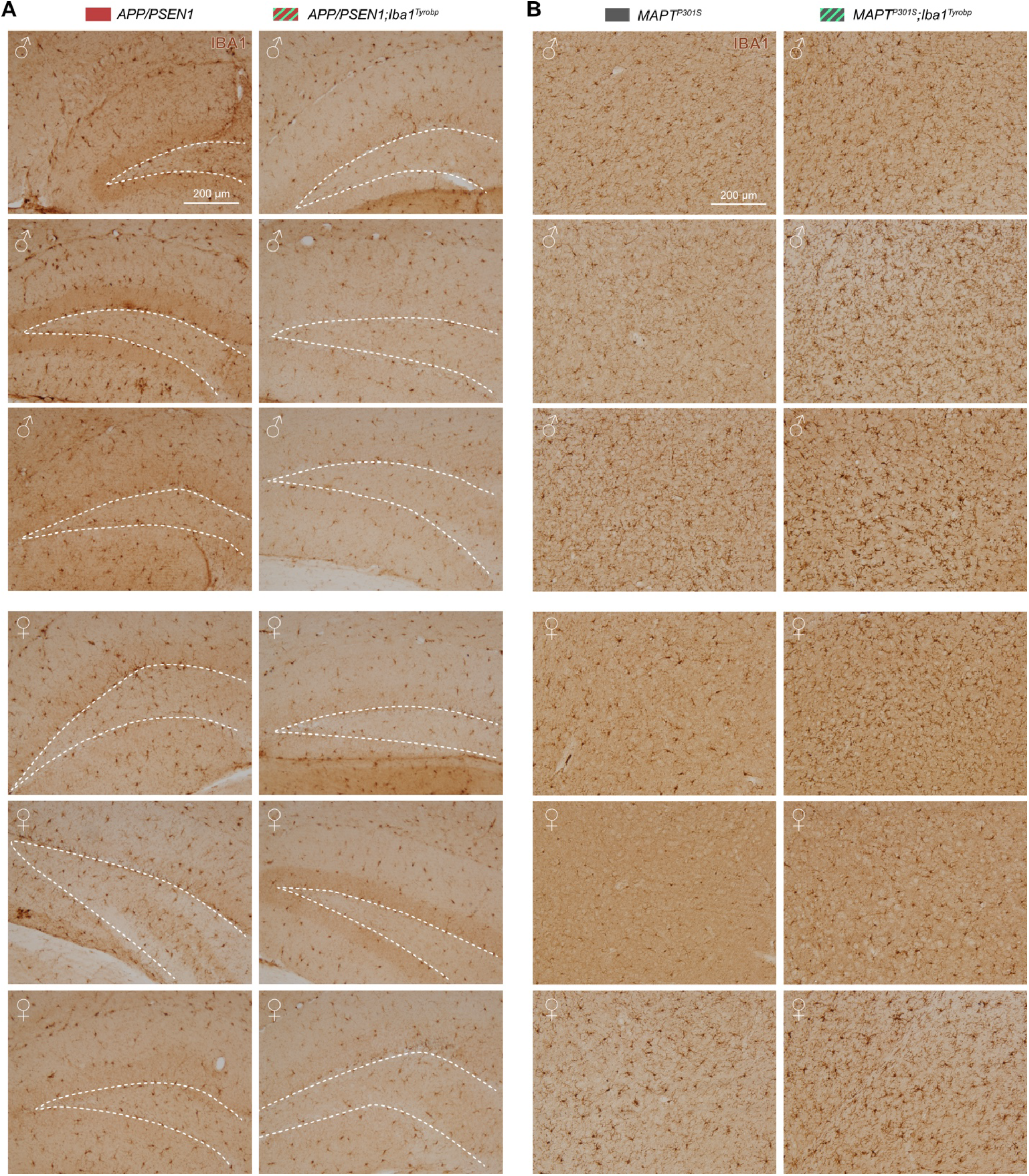
Anti-IBA1 immunohistochemistry in 4-month-old *APP/PSEN1* and *MAPT^P301S^* mice with and without the *Iba1^Tyrobp^* transgene. (A) DAB-immunohistochemistry with anti-IBA1 in 4-month-old *APP/PSEN1* and *APP/PSEN1;Iba1^Tyrobp^* mice. (B) DAB-immunohistochemistry with anti-IBA1 in 4-month-old *MAPT^P301S^* and *MAPT^P301S^;Iba1^Tyrobp^* mice. N = 3 males and 3 females per genotype. Scale bar = 200μm.

**Supplementary Figure 3:**
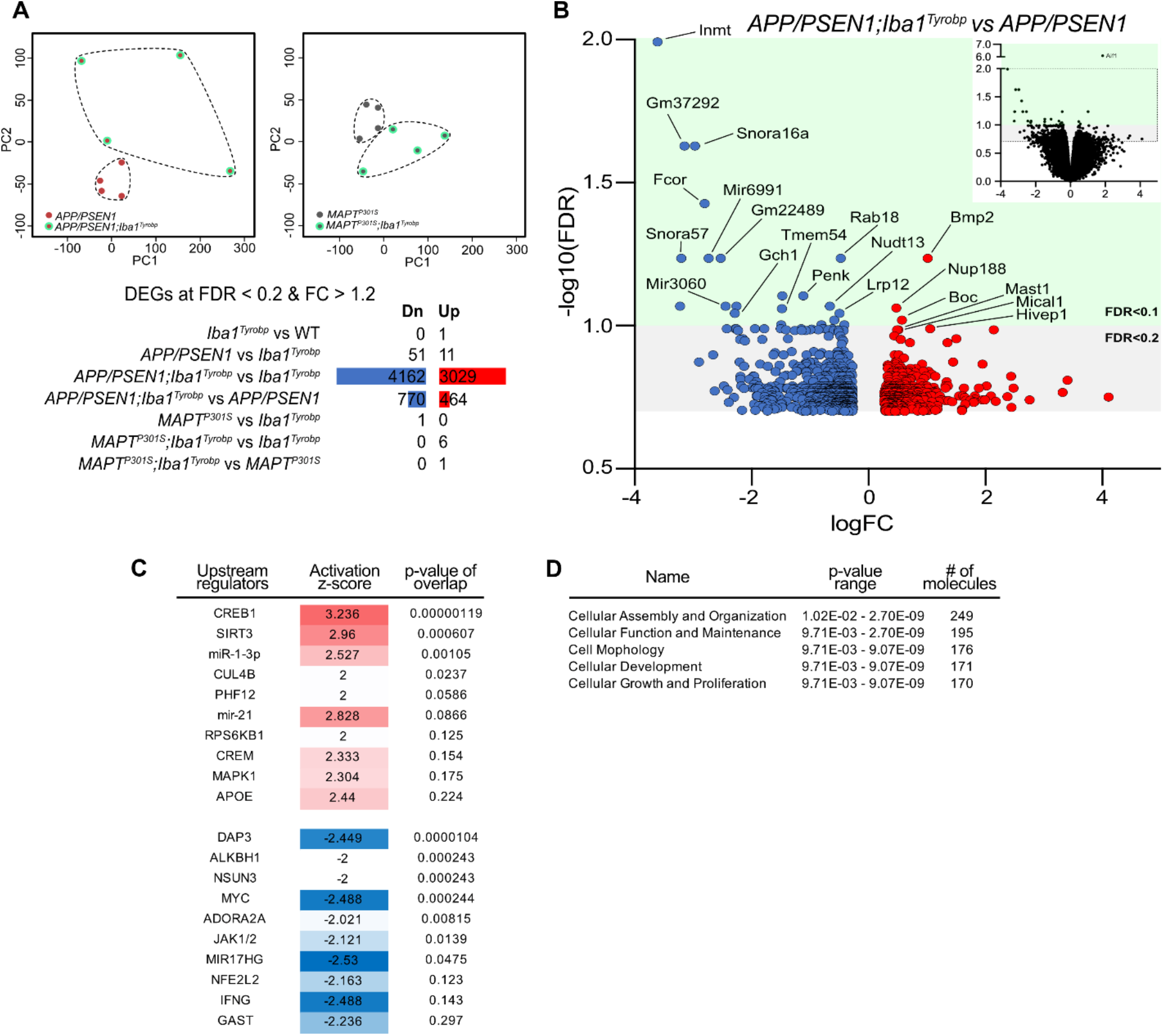
Bulk RNA sequencing analysis of *APP/PSEN1* or *MAPT^P301S^* on *Iba1^Tyrobp^* background. (A) RNA sequencing was performed on hippocampi from 4-month-old male WT, *Iba1^Tyrobp^, APP/PSEN1, APP/PSEN1;Iba1^Tyrobp^, MAPT^P301S^* or *MAPT^P301S^;Iba1^Tyrobp^*mice, N = 4/genotype. Top: Principal Component Analysis (PCA) of the *APP/PSEN1* vs *APP/PSEN1;Iba1^Tyrobp^* RNA sequencing samples (left) and *MAPT^P301S^* vs *MAPT^P301S^;Iba1^Tyrobp^* RNA sequencing samples (right). Bottom: Numbers of DEGs (FDR <0.2 and FC > 1.2) in the genotype comparisons. Blue and red bars represent the number of significantly down- and up-regulated genes, respectively. (B) Volcano plot representation of the DEGs (FDR <0.2 and FC > 1.2) from the *APP/PSEN1;Iba1^Tyrobp^* vs *APP/PSEN1* comparison. The top right quadrant represents the DEGs that have been graphed (all genes with FDR < 0.2 except *Aif1*). (C-D) Ingenuity Pathway Analysis (IPA) was used to identify the top upstream regulators and predicted activation states (C) as well as the top molecular and cellular functions (D) in the *APP/PSEN1;Iba1^Tyrobp^* vs *APP/PSEN1* comparison.

**Supplementary Figure 4:**
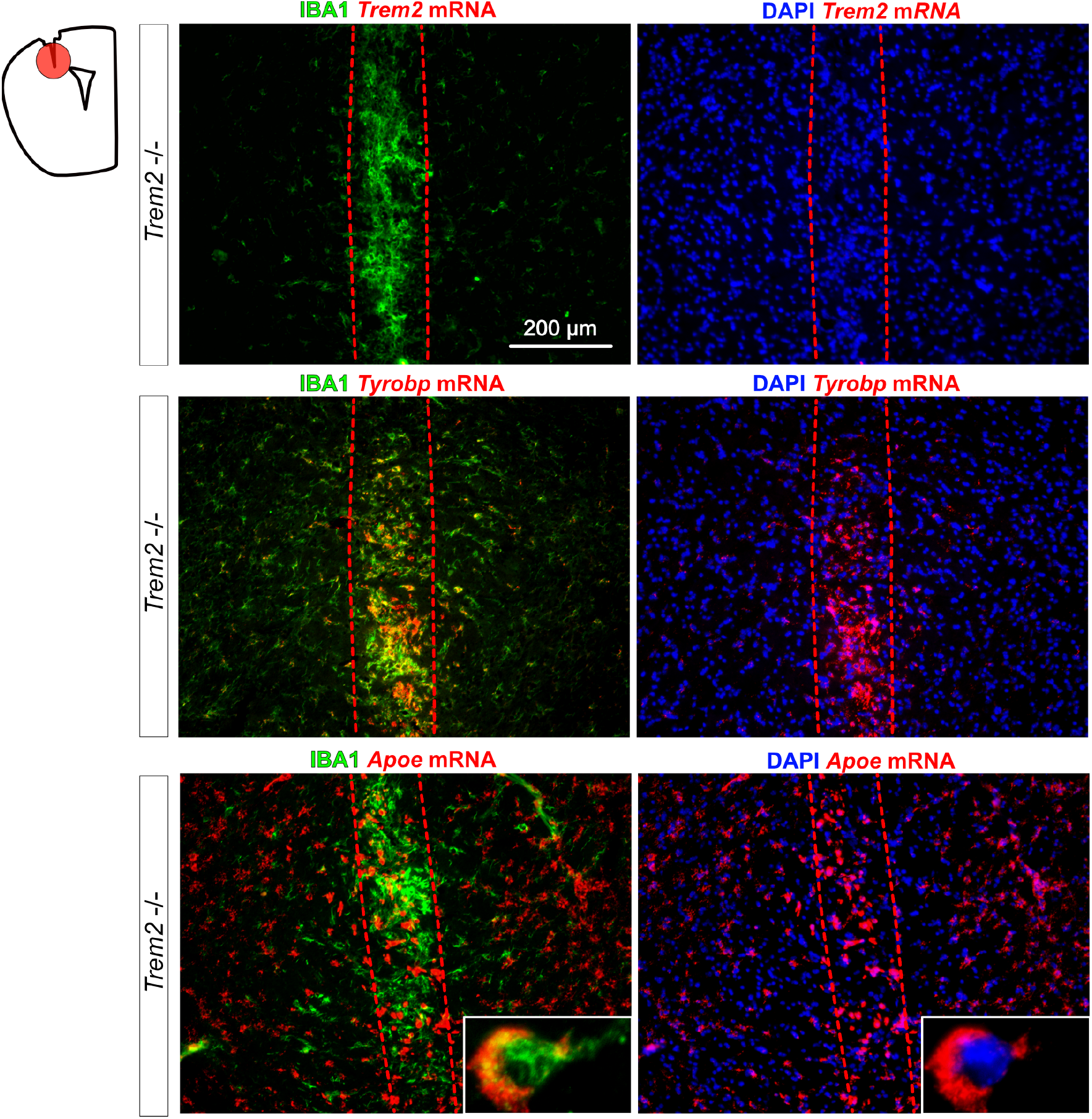
*Tyrobp* and *Apoe* mRNAs are increased in microglia recruited around the needle tract of stab injured-*Trem2^−/−^* mice. RNA fluorescent *in situ* hybridization for *Trem2* or *Tyrobp* or *Apoe* mRNA double-labeled with an anti-IBA1 antibody (green) or DAPI (blue) in *Trem2^−/−^* mice injured as described. The red dot line indicates the needle track. Scale bar = 200 μm.

